# How individual differences shape ERP responses to visual statistical learning

**DOI:** 10.1101/2025.07.30.667804

**Authors:** Joanna Morris, Emma Kealey, Frankie Greene, Joemari Pulido, Tess Rooney, Natalia Alzate, Ethan Moore, Crismar Ramos-Marte

## Abstract

Statistical learning (SL) enables the extraction of regularities from sensory input, yet the neural dynamics supporting this process—particularly in the visual modality—remain incompletely understood. Sixty-seven adults were familiarized with a continuous stream of shape sequences containing statistical structure that defined shape triplets. We recorded EEG to familiar sequences (presented in isolation) and unfamiliar foils. Both early (N100) and late (N400) event related potential (ERP) components were significantly more negative for unfamiliar than familiar sequences, reflecting robust neural sensitivity to learned structure. Notably, these familiarity effects were evident in both high- and low-performing participants and were not predicted by overall behavioral sensitivity, suggesting that neural indices of learning can emerge independently of explicit recognition. Follow-up analyses incorporating trial-level accuracy revealed a striking crossover interaction: for sensitive participants, ERP familiarity effects were stronger on correct trials, whereas for insensitive participants, effects were larger on incorrect trials. These findings highlight a dissociation between neural and behavioral measures of statistical learning and underscore the value of ERPs in capturing latent learning processes that may elude conscious awareness.

## Introduction

Our perceptual, cognitive, and social environments are structured by statistical regularities. Statisical learning (SL) describes the set of processes by which we detect and internalize these patterns. Frost et al. define SL as “perceiving and learning any forms of patterning in the environment that are either spatial or temporal in nature” and subsequently using that statistical information to construct mental representations (p. 1130) [1]. SL has garnered sustained interest in cognitive neuroscience because many core cognitive functions—such as contextual cueing, scene perception, visual word recognition, and face perception—depend on the ability to learn structured patterns in sensory input [1–4]. Reading acquisition, for instance, requires sensitivity to a variety of regularities: the mappings between letters or letter clusters and corresponding sounds; associations between morphological elements such as prefixes and suffixes and their semantic content; the spatial arrangement of graphemes and their relation to lexical status; the probabilistic co-occurrence of letters within words and of words within sentences; and correlations between orthographic patterns and syntactic roles [1].

Given SL’s foundational role in perceptual and cognitive processes, researchers have increasingly examined whether individual differences in reading and language abilities reflect underlying variability in SL capacity. To date, the majority of this work has relied exclusively on behavioral measures. Although informative, behavioral tasks provide only a limited view of the learning process. They typically capture the final outcome of a complex sequence involving perceptual analysis, cognitive evaluation, response selection, and motor execution. Moreover, behavioral assessments often require participants to make explicit judgments about what they have learned, despite the fact that SL is believed to operate largely implicitly, outside of conscious awareness. This mismatch between the implicit nature of SL and the explicit demands of typical behavioral tasks may lead to an underestimation of learning effects.

To address these limitations, researchers have turned to event-related potentials (ERPs) as a complementary method for studying SL. ERPs offer high temporal resolution and are time-locked to stimulus onset, enabling fine-grained analyses of when learning-related changes occur during information processing. Their capacity to differentiate components based on latency and scalp topography also makes them well-suited for isolating distinct neurocognitive mechanisms [5]. These methodological advantages position ERPs as a powerful tool for investigating how statistical regularities are detected and internalized, particularly when such learning unfolds implicitly.

The overarching goal of this study is to better understand how individual differences in sensitivity to statistical structure are reflected in the brain, with a specific focus on visual SL. We examine this question using ERPs, which provide a window into the time course and neural mechanisms of learning. In what follows, we first introduce the two ERP components of interest—the N100 and N400—and review their relevance to SL. We then summarize findings from ERP studies of auditory SL, before turning to visual SL, highlighting discrepancies in the literature and identifying key open questions that motivate the present study. Finally, we describe the design and aims of the current study, outlining our specific predictions and how they address gaps in prior work.

### The N400 and N100 Components

Here, we use the hybrid term *N1/N100* to refer to the early negative-going ERP component that typically peaks around 100 ms post-stimulus and is commonly labeled *N1* in visual paradigms and *N100* in auditory paradigms. Throughout the text, we retain the original terminology used in each cited study but interpret these labels as referring to functionally equivalent components.

The N1/N100 and N400 are two event-related potential (ERP) components commonly studied in relation to perceptual and language processing. The N1/N100 is a negative-going, sensory-evoked component that peaks between approximately 80 and 120 ms after stimulus onset. It is typically distributed over fronto-central regions in the auditory modality and posterior-lateral regions in the visual modality. The N1/N100 is thought to reflect early perceptual encoding of physical stimulus features such as pitch, location, and shape [6, 7]. This component is known to be modulated by stimulus salience, novelty, and particularly attention: attended stimuli generally elicit larger N1/N100 amplitudes than unattended ones [7, 8]. These characteristics suggest that the N1/N100 reflects an early stage of information processing that is sensitive to both bottom-up and top-down influences.

The N400, by contrast, is a negative-going component that typically peaks between 300 and 500 ms post-stimulus and is maximal over centro-parietal sites. Originally identified in studies of language, the N400 is traditionally associated with semantic processing, particularly the integration of a word into its sentence context [9]. However, later research has expanded its interpretation to include sensitivity to predictability, associative relationships, and the ease of integrating a stimulus into a broader mental context [10]. In this view, the N400 reflects a dynamic updating process in which incoming input is evaluated against prior expectations or structured representations. Unlike the N1/N100, which reflects early perceptual and attentional processes, the N400 is more closely tied to higher-order meaning-based or expectancy-driven computations.

### ERP Studies of Auditory Statistical Learning

In auditory SL paradigms, participants are exposed to continuous streams of syllables or tones, within which “words” are defined by high transitional probabilities (TPs). A core aim of ERP research in this domain has been to identify neural signatures of segmentation and learning as participants listen to these structured sequences. Two components have received particular attention: the N400, which reflects integrative processing and representational coherence, and the N100, associated with early perceptual segmentation and attentional orienting.

#### The N400: Late Integration and the Formation of Structured Representations

The N400 has been consistently observed across auditory statistical learning (SL) studies and is widely interpreted as reflecting the formation of emergent lexical-like representations—such as syllable triplets—based on distributional regularities in the input. In early work, [11] reported that N400 amplitude increased following exposure to statistically coherent syllable streams, even among low-performing learners who did not exhibit neural evidence of early segmentation processes (i.e., they lacked earlier ERP effects typically associated with the detection of transitional probabilities). This finding suggests that the N400 may reflect the formation of protolexical representations independent of early perceptual segmentation.

Building on this early work, Abla et al. [12] found that highly successful learners exhibited enhanced N400 amplitudes at word onsets early in training, followed by decreasing amplitude across subsequent blocks—consistent with the development of familiarity-based lexical representations. Moderately successful learners showed a similar pattern, but it emerged later in training, suggesting a more gradual consolidation process.

More recently, Soares et al. [13, 14] extended these findings by manipulating both the strength of transitional probabilities (TPs) between and within statistically defined units and the nature of task instructions, which either directed participants’ attention to the embedded structure (explicit learning) or did not (implicit learning). In Soares et al. [13], the N400 was larger for high-TP (easier) pseudowords than for low-TP (harder) ones, regardless of whether participants received implicit or explicit instructions. In a follow-up study, Soares et al. [14] found that N400 amplitude for high-TP pseudowords increased over time, particularly under explicit learning conditions and in the later phases of the task.

Together, these results support the view that the N400 reflects the formation and consolidation of structured mental representations. They also suggest that this component is a relatively robust neural index of statistical learning, emerging across a range of task demands and levels of behavioral performance.

#### The N100: Early Segmentation Driven by Prosodic and Statistical Cues

While the N400 has been robustly observed across auditory statistical learning studies, the N100 has appeared more inconsistently and primarily under specific conditions that facilitate perceptual segmentation. In particular, studies that provide additional cues to boundary structure—such as prosodic stress, high transitional probabilities, or explicit task instructions—are more likely to elicit early N100 effects.

Prosodic cues appear to play a critical role. In a naturalistic speech paradigm, Sanders and Neville [15] found that word-initial syllables elicited larger N100 responses than medial ones and that stressed syllables produced stronger effects, indicating that prosodic emphasis can enhance early auditory processing. Similarly, Cunillera et al. [16] manipulated syllabic stress and found that N1 and N400 effects emerged only in the presence of stress cues. Stressed word onsets elicited larger N1 amplitudes than nonwords, and N400 amplitudes were enhanced for words versus nonwords in the stressed condition, suggesting that prosody facilitates both boundary detection and lexical-like integration.

In the absence of prosodic cues, early segmentation effects have been observed only when transitional probability structure was especially strong or explicitly highlighted. For example, in Soares et al. [13], N100 amplitude was modulated by a three-way interaction between word type (high vs. low TP), task instruction (implicit vs. explicit), and exposure block. Effects were strongest for high-TP pseudowords under explicit instructions in the later blocks of the task. A follow-up study, Soares et al. [14], replicated and extended these findings, again showing that both N100 and N400 amplitudes were larger for high-TP words, particularly in the second half of the task and under explicit learning conditions. These results suggest that both attention and TP structure are critical in modulating early sensory responses, and that N100 enhancement reflects the initial detection of coherent structure in the input.

Other studies have found that N100 effects emerge only after training or with extended exposure, particularly among high-performing learners. In a foundational auditory SL study, Sanders et al. [11] showed that N100 amplitudes increased for word onsets—but only after training and only among participants who successfully learned the embedded structure. Likewise, Abla et al. [12] found that high learners exhibited enhanced N100s to word-onset tones in the first exposure block, while middle learners showed similar effects only by the third block. Low learners showed no N100 modulation. Notably, these N100 effects were positively correlated with transitional probability, supporting the idea that the N100 indexes early segmentation based on learned statistical structure.

In contrast, when none of these cues or conditions are present, the N100 often fails to emerge. For example, in a follow-up to their stress-manipulation study, Cunillera et al. [17] found no N100 difference between structured and random streams, even across multiple exposure blocks. The same study revealed a robust N400 effect that followed an inverted U-shaped trajectory, peaking in the second block and declining thereafter. Similarly, Abla and Okanoya [18] reported no N100 modulation in a visual SL task despite observing clear N400 effects. Together, these findings suggest that early segmentation responses are contingent on conditions that guide attention to statistical structure, whereas the N400 may reflect the integration of structure even in the absence of such cues.

These patterns also help clarify a point raised earlier: low-performing learners who fail to exhibit early ERP effects typically associated with the detection of transitional probabilities nonetheless often show N400 modulation. This dissociation underscores the idea that the N100 and N400 index distinct stages of learning. While the N100 reflects perceptual segmentation mechanisms, often driven by attentional support or cue salience, the N400 reflects post-segmentational integration or the formation of protolexical representations, which may proceed even in the absence of overt boundary detection.

Taken together, these findings support a temporal cascade model of statistical learning. The N100 appears to mark the initial parsing of sensory input into candidate units, contingent on attention, cue availability, or learning proficiency. The N400, by contrast, reflects the subsequent integration of these units into more stable representations, indexing familiarity, coherence, and prediction. Importantly, while both components are modulated by statistical learning, the N400 emerges more consistently, highlighting its central role in tracking the formation of statistically-defined units—even when earlier segmentation processes are weak or undetected.

### ERP Studies of Visual Statistical Learning

While most ERP research on statistical learning (SL) has focused on the auditory domain, statistical regularities are also readily extracted from visual input. Indeed, a central claim in the statistical learning literature is that these mechanisms are largely domain general, enabling the extraction of regularities across different sensory modalities, although the exact nature and limits of these mechanisms remain active topics of investigation. However, ERP studies of visual SL have been less common and typically employ different task designs than their auditory counterparts. As a result, our understanding of the neural dynamics of visual SL, and how they compare to those observed in auditory studies, remains limited. In particular, whereas auditory studies often use continuous streams of syllables or tones to examine segmentation processes, visual studies have tended to use target-detection paradigms that prioritize statistical contingency over sequence learning.

Two prominent studies of visual statistical learning, Jost et al. [19] and Daltrozzo et al. [20], used paradigms based on pairwise contingencies between predictor and target stimuli. Both identified late positive ERP components (e.g., P300/P600) associated with the learning of high-probability associations over time. While these studies support the domain-generality and developmental robustness of SL, their designs do not involve continuous streams or structured sequences. As such, they are not well suited to addressing questions about sequence segmentation or the role of early components like the N100 and N400.

Only one study to date, that of Abla and Okanoya [18], has applied a continuous stream paradigm in the visual domain that closely mirrors those used in auditory SL research. In this study, participants viewed sequences of geometric shapes organized into repeating triplets and were later tested on their ability to discriminate familiar from unfamiliar sequences. ERP responses were recorded during exposure to the continuous stream. As in the corresponding auditory study by Abla et al. [12], the focus was on shape triplet onsets as potential segmentation points.

Notably, Abla and Okanoya observed an N400 effect at triplet boundaries: high learners showed significantly larger N400 amplitudes for the first shape of each triplet compared to medial or final shapes. Although the authors interpreted this as reflecting predictability-based processing or online segmentation, we adopt the view that the N400 reflects post-segmentational integration or the formation of protolexical representations, in line with interpretations from auditory SL research. The N400 was largest early in learning and declined across exposure blocks, suggesting that as the statistical structure became internalized, demands on integrative processes lessened. In contrast to many auditory SL studies, which have reported N400 effects even in low-performing learners, low performers in this visual task showed no significant N400 modulation, though the authors noted a numerical trend that might have reached significance with extended exposure. This pattern suggests that the integrative processes indexed by the N400 may emerge more slowly, or require stronger learning—in the visual modality. Nonetheless, the findings extend the core auditory result to visual sequences, supporting the domain-generality of later-stage integration mechanisms in statistical learning.

Strikingly, however, Abla and Okanoya did not observe any modulation of the N100 component, even among high-performing learners. This stands in contrast to their earlier auditory study, which found robust N100 enhancement for high learners at triplet onsets. The absence of an N100 effect in the visual version of the task raises a key question: Does visual statistical learning engage early perceptual segmentation mechanisms to the same extent as auditory learning?

This question becomes particularly salient when considered alongside auditory studies that failed to elicit N100 effects under certain conditions. As discussed earlier, the presence of an N100 has been shown to depend on factors such as prosodic cues [15, 16], extremely high transition probabilities [13, 14], and explicit instruction directing attention to the structure. When such cues are absent, as in the visual paradigm of Abla and Okanoya [18], early segmentation effects indexed by the N100 may fail to emerge, even when later-stage processing (reflected in the N400) remains intact.

A second, and equally notable, departure from the auditory SL literature is the absence of N400 effects in low-performing learners. In auditory studies, the N400 has often been observed even in participants who show little or no behavioral evidence of learning, suggesting that post-segmentational integration can occur independently of overt task performance. In contrast, Abla and Okanoya found that only high learners exhibited N400 modulation at triplet boundaries; low learners showed no significant N400 effect, despite numerical trends in the expected direction. This pattern suggests that the integrative processes indexed by the N400 may require stronger or more sustained learning in the visual modality than in the auditory domain.

In short, while both modalities support statistical learning and show N400 effects associated with representational integration, two important discrepancies emerge: the absence of early N100 modulation in the visual task and the restriction of N400 effects to high-performing learners. The former may reflect differences in the way visual and auditory systems support perceptual segmentation. The latter may suggest that neural indices of integrative processing are less likely to arise in the absence of successful learning in the visual modality. Together, these findings raise the possibility that visual statistical learning relies on different, or less automatic, mechanisms than its auditory counterpart. They may also reflect limitations in the temporal resolution of the visual system for parsing continuous input or reduced engagement of early perceptual circuits in the absence of spatial or prosodic segmentation cues.

TThese differences raise the possibility that there are modality-specific constraints on statistical learning. They also motivate the use of ERP methods to test whether neural markers of learning observed in auditory paradigms—such as the N100 and N400—extend to the visual domain. Clarifying when early segmentation and later integration processes are engaged during visual SL is essential for understanding how statistical structure is extracted and represented in the brain.

### The current study

This study aimed to advance our understanding of individual differences in visual statistical learning (visual SL) by examining how sensitivity to statistical structure is reflected in neural activity. We used event-related potentials (ERPs) to track learning-related brain responses, focusing on two components—the N100 and N400—that have previously been linked to the processing of statistical regularities in sensory input.

Prior work on auditory SL has consistently implicated both components in learning: the N100 as an index of early perceptual segmentation and the N400 as a marker of post-segmentational integration or representational coherence. In the only prior study using a continuous visual triplet paradigm comparable to auditory designs, Abla and Okanoya [18] observed an N400 effect in high-performing learners but found no evidence of N100 modulation—even among those who demonstrated behavioral learning. This pattern raises two important open questions: First, does visual SL reliably engage early segmentation mechanisms, or is the N100 component modality-specific or cue-dependent? Second, can neural evidence of learning—as indexed by the N400—be observed in individuals who fail to demonstrate behavioral sensitivity to statistical structure, as has been reported in auditory studies?

To address these questions, we tested 67 adult participants using a visual SL paradigm closely modeled on that of Abla and Okanoya [18]. During the familiarization phase, participants were exposed to continuous sequences of shape triplets without explicit segmentation cues. In a subsequent test phase, we recorded ERPs while participants viewed familiar and unfamiliar triplets and made two-alternative forced-choice decisions. Participants were then categorized as sensitive (above-chance performance) or insensitive (chance-level or below) based on their test accuracy. This design allowed us to examine how the N100 and N400 components varied as a function of both familiarity (familiar vs. unfamiliar sequences) and sensitivity (individual learning performance).

Importantly, our sample size (N = 67) afforded substantially greater statistical power than previous ERP studies of visual SL, such as Abla and Okanoya [18], which included only 18 participants. The increased power allowed us to test for the reliability of both early and late ERP effects across a wide spectrum of learning sensitivity, enabling more precise comparisons between high- and low-performing participants. The study was designed to address three key questions: (1) Do the N100 and N400 components reliably differentiate familiar from unfamiliar shape sequences, indicating neural sensitivity to statistical structure? (2) Are these ERP effects modulated by individual differences in behavioral sensitivity? (3) Can neural evidence of learning emerge even in the absence of behavioral learning, particularly for later components like the N400?

Based on prior auditory SL research, we predicted that the N400 would be enhanced for unfamiliar sequences relative to familiar ones, particularly among sensitive learners, reflecting increased effort at post-segmentational integration. We also hypothesized that if early segmentation mechanisms are engaged in the visual modality, then N100 amplitude should differentiate familiar from unfamiliar sequences as well. However, given the lack of N100 effects in previous visual SL studies, we considered this prediction exploratory and expected any such effects to be more contingent on individual sensitivity and task demands.

## Materials and methods

### Participants

We tested 76 participants recruited from the UMass–Five Colleges consortium community in Amherst, MA. Participants were included if they were between the ages of 18 and 26, right-handed, reported having normal or corrected-to-normal vision with no history of cognitive or reading-related difficulties and were native English speakers with no other languages spoken in the home before the age of 3. Three participants were excluded for exceeding the protocol’s age range, and six were excluded due to data loss, resulting in a final sample of 67 participants (age range: 18–25 years, M = 19.6). Of these, 49 identified as female, 17 as male, and one did not report their sex. All participants provided written informed consent prior to the start of the study and were paid for their participation. All materials and procedures were reviewed and approved by the IRB committee at Hampshire College.

To assess whether our study was adequately powered to detect effects of similar magnitude to those reported in prior ERP studies of SL, we conducted a post hoc power analysis using effect sizes from the literature. Reported *η*^2^ values for key effects ranged from .25 to .44. Using these values to estimate Cohen’s *f* and applying them to our sample size (*n* = 67), we found that our study had high power (*≥* .95) to detect effects of similar size.

### EEG Data Acquisition and Processing

EEG activity was recorded at 500 Hz from 31 Ag/AgCl electrodes with integrated impedance conversion (active electrodes) placed according to the extended 10–20 system and attached to an elastic cap (actiCAP, BrainProducts GmbH, Gilching, Germany). We used an initial reference-free montage during data acquisition, with data later re-referenced offline to the algebraic average of the left and right mastoid. The ground electrode was placed at Fpz. Impedances were kept below 25 *k*Ω, which is within the acceptable range for active electrodes. Signals were amplified using the actiCHamp Plus EEG system (BrainProducts GmbH, Gilching, Germany) and recorded with the BrainVision Recorder software (BrainProducts GmbH, Gilching, Germany). Vertical eye movements and blinks were monitored using electrode sites FP1 and FP2.

The data were preprocessed using EEGLAB [21] and ERPLAB [22] prior to statistical analysis. The data were filtered with a bandpass filter of 0.1–30 Hz (zero-phase shift Butterworth), downsampled to 200 Hz, and re-referenced to the algebraic average of the left and right mastoid. Independent component analysis (ICA) was performed to remove ocular and muscular artifacts. Components reflecting eye blinks and saccades were identified based on their characteristic topography and time course and were removed from the data before further processing. Bad electrodes were identified and interpolated using spherical interpolation.

The data were segmented into epochs extending from –200 ms to 1000 ms relative to the onset of each shape. Baseline correction was applied prior to artifact rejection, using the 100 ms pre-stimulus interval (–100 to 0 ms relative to stimulus onset). This baseline period is commonly used in ERP research to normalize voltage fluctuations and isolate stimulus-evoked activity. Epochs containing amplitudes exceeding *±*100 *µ*V were excluded. After artifact rejection, 96.5% of trials were retained.

The epoched data were averaged to create event-related potentials. Mean amplitudes were extracted from two post-stimulus intervals (75–175 ms for the N100 and 350–500 ms for the N400), based on prior literature defining these components in visual word recognition paradigms.

### Sensitivity Classification

We assessed participants’ ability to discriminate between familiar and unfamiliar sequences during the testing phase based on their accuracy. Accuracy was defined as the proportion of trials on which participants correctly identified a familiar triplet as familiar or an unfamiliar triplet as unfamiliar. We then calculated a *d*′ score for each participant as a measure of sensitivity to the statistical structure of the triplets (*d*′ = *z*(*p*_correct|familiar_) *− z*(*p*_incorrect|unfamiliar_)). Participants with *d*′ *>* 0 were classified as sensitive, indicating above-chance discrimination performance, whereas those with *d*′ *≤* 0 were classified as insensitive. Based on this criterion, 51 participants (76.1%) were categorized as insensitive and 17 (25.4%) as sensitive. The mean accuracy score for sensitive participants was 79.3%, compared to 51.7% for insensitive participants.

### Statistical Analysis

Statistical analyses were conducted using linear mixed-effects modeling in R version [23] within RStudio [24]utilizing the *afex* package [25]. A mixed-effects approach was selected over traditional repeated-measures ANOVA because it accommodates unbalanced data, accounts for subject-level variability, and does not require sphericity assumptions, making it well suited to ERP data with multiple repeated measurements per participant [26–28].

The primary factors of interest were *Sensitivity* (Sensitive, Insensitive; between-subjects) and *Familiarity* (Familiar, Unfamiliar; within-subjects). Their interaction was included to test whether the effect of Familiarity varied as a function of individual differences in sensitivity to statistical structure. ERP amplitude was modeled without additional fixed effects for scalp location (e.g., Anteriority or Laterality), as these spatial factors were not central to our hypotheses and including them would substantially increase the number of statistical comparisons, inflating the Type I error rate [29]. Instead, to account for multiple spatially correlated observations, spatial variance was treated as a random factor by modeling electrode site as a random intercept nested within participant. The final model was specified as:

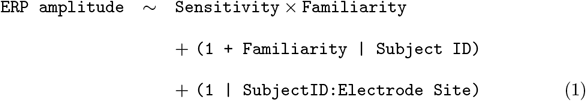

Random intercepts for each participant captured baseline variability in ERP amplitude, while random slopes for Familiarity allowed for individual differences in the size and direction of familiarity effects. Electrode sites were nested within participants to retain spatial variability while avoiding the multiple comparisons and inflated familywise error rate associated with modeling scalp region as a fixed effect [29]. This approach also reduces the number of hypothesis tests and increases statistical power relative to models that treat Anteriority and Laterality as fixed factors. Finally, it avoids the information loss and underestimated standard errors associated with averaging across electrodes, and it more accurately reflects the nested structure of ERP data than a cross-classified model treating subject and electrode as independent random effects.

Models were estimated using restricted maximum likelihood (REML), and the Kenward-Roger approximation was used to calculate denominator degrees of freedom and improve control of Type I error [30]. Fixed effects were tested using Type III Wald F tests. Estimated marginal means (EMMs), standard errors (SE), and 95% confidence intervals (CI) were obtained using the emmeans package [31], and pairwise comparisons were Bonferroni-corrected to account for multiple testing. The full random effects structure was retained following recommendations for confirmatory hypothesis testing in ERP designs [27].

## Results

To address our research questions, we conducted a series of statistical analyses focused on two ERP components—the N100 and N400—that are commonly associated with early perceptual and later semantic or integrative processing, respectively. The primary aim was to determine (1) whether these components reliably differentiate statistically familiar from unfamiliar shape sequences, and (2) whether neural sensitivity to structure varies with individual differences in behavioral performance. We begin by examining the effects of our manipulated variable, Sequence Familiarity (familiar vs. unfamiliar sequences), and the individual difference factor Sensitivity to Statistical Structure (sensitive vs. insensitive participants), focusing first on ERP responses in the N400 time window. Based on prior research, we predicted that: (1) familiar sequences would elicit more negative N400 responses than unfamiliar sequences; (2) sensitive participants would show more negative N400 responses than insensitive participants; and (3) the difference between familiar and unfamiliar sequences would be greater for sensitive than for insensitive participants, resulting in an interaction between familiarity and sensitivity. Given prior evidence that the N400 is a robust marker of SL, we expected these effects to emerge most clearly in this component. However, if the absence of reliable N100 effects in earlier visual studies was due to limited statistical power rather than a true absence of signal, we would expect that a similar pattern of effects may also be observed in the N100 time window. Following these primary analyses, we conduct a second set of models incorporating behavioral Accuracy, to assess whether accuracy interacts with Familiarity or Sensitivity, and to explore whether ERP responses reflect learning even in the absence of overtly correct performance.

### Model with Familiarity and Sensitivity

We fit separate linear mixed-effects models for each component (N100, N400), using an identical structure. Fixed effects included Sensitivity (Sensitive, Insensitive), Familiarity (Familiar, Unfamiliar), and their interaction. Random effects included by-subject intercepts and Familiarity slopes, along with electrode-level intercepts nested within subjects. Likelihood ratio tests confirmed that including both the random Familiarity slope, *χ*^2^(2) = 516.42–562.11, *p <* .001, and the nested electrode intercept, *χ*^2^(1) = 158.85–165.97, *p <* .001, significantly improved model fit for both components. The N100 model revealed a significant main effect of Familiarity, *F* (1, 65) = 27.44, *p <* .001, 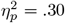, with larger (i.e., more negative) amplitudes for familiar compared to unfamiliar sequences (*M* = *−*3.55 *µ*V, *SE* = 0.51, 95% CI [–4.56, –2.54] vs. *−*0.90 *µ*V, *SE* = 0.26, 95% CI [–1.42, –0.37]), *d* = *−*0.55, 95% CI [–0.63, –0.47]. The main effect of Sensitivity was not significant, *F* (1, 65) = 2.50, *p* = .119, *η*^2^ = .04, although amplitudes for sensitive participants were somewhat more negative than those of insensitive participants (*M* = *−*2.72 *µ*V, *SE* = 0.54, 95% CI [–3.79, –1.64] vs. *−*1.73 *µ*V, *SE* = 0.31, 95% CI [–2.36, –1.10]), *d* = *−*0.22, 95% CI [–0.31, –0.13]. The Sensitivity *×* Familiarity interaction was not significant, *F* (1, 65) = 1.03, *p* = .313,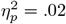. The model explained 8.0% of the variance via fixed effects 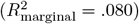 and 57.2% overall 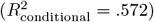. (See Fig 1).

**Fig 1.**
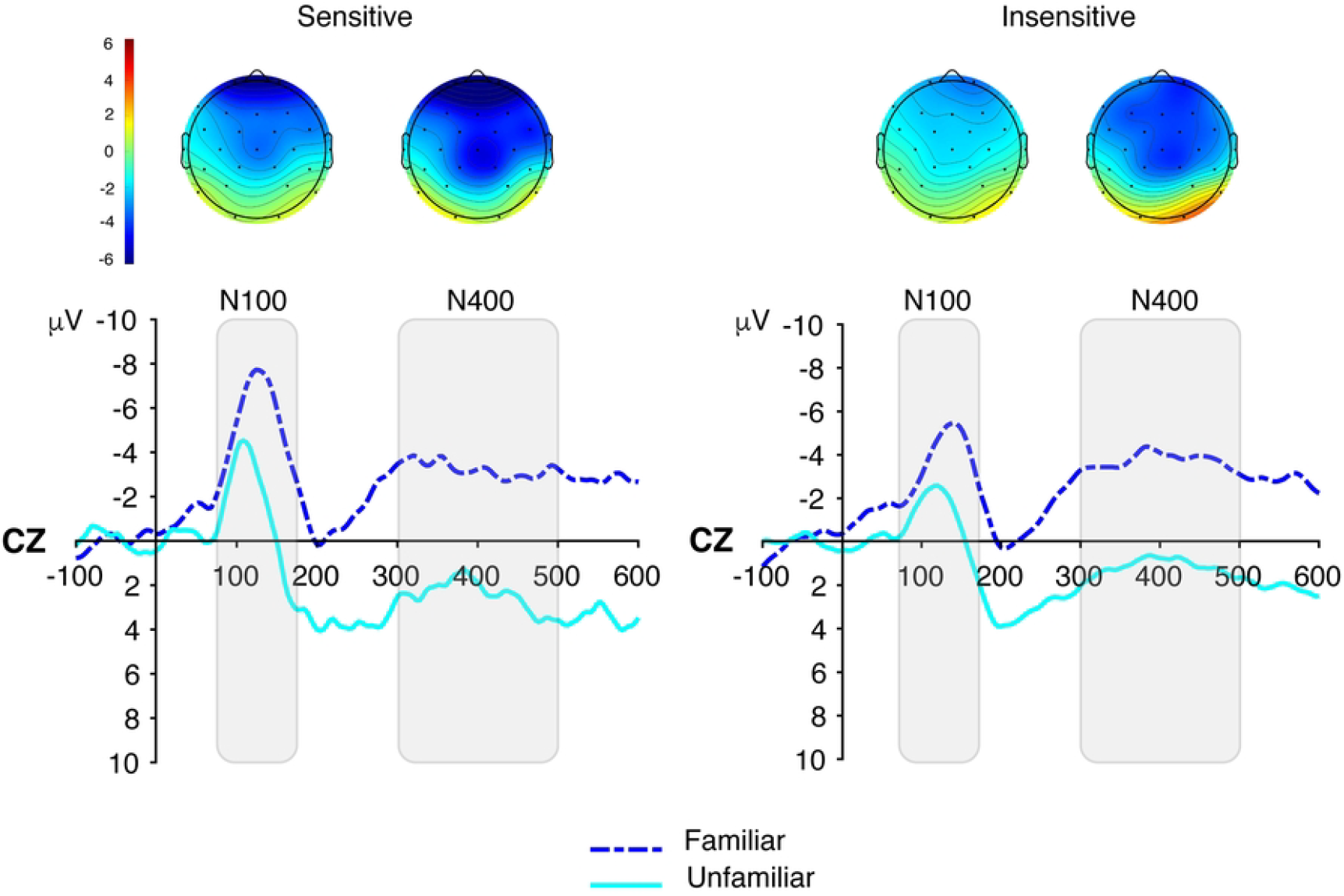
ERP results illustrating the effects of familiarity on N100 and N400 amplitudes for sensitive and insensitive participants. A: Scalp topographies showing the distribution of mean amplitude in the N100 (75–175 ms) and N400 (300–500 ms) time windows. B: Grand-average ERP waveforms at electrode Cz, time-locked to the onset of shape triplets, plotted separately for Familiar and Unfamiliar Triplets. Negative values are plotted up.

### ERP responses to Familiarity by learner group

To further examine whether neural responses to statistically familiar sequences differed by learning sensitivity, we conducted separate linear mixed-effects models for sensitive and insensitive participants, with Familiarity as a fixed effect and by-subject random slopes. These analyses were performed separately for the N100 and N400 components.

For sensitive participants, Familiarity significantly predicted N100 amplitude, *F* (1, 16) = 9.44, *p* = .007, with a large effect size, 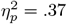, 95% CI [0.08, 1.00]. The N100 was more negative for familiar sequences (*M* = *−*4.30, *SE* = 1.01, 95% CI [*−*6.44, *−*2.17]) than for unfamiliar sequences (*M* = *−*1.13, *SE* = 0.53, 95% CI [*−*2.26, 0.00]), *t*(16) = *−*3.07, *p* = .007, difference = *−*3.17, *SE* = 1.03. For insensitive participants, Familiarity also significantly predicted N100 amplitude, *F* (1, 49) = 20.11, *p <* .001, with a large effect size, 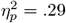, 95% CI [0.13, 1.00]. N100 amplitude was more negative for familiar items (*M* = *−*2.80, *SE* = 0.48, 95% CI [*−*3.77, *−*1.84]) than for unfamiliar items (*M* = *−*0.66, *SE* = 0.25, 95% CI [*−*1.15, *−*0.17]), *t*(49) = *−*4.48, *p <* .001, difference = *−*2.14, *SE* = 0.48.

In the N400 data, For sensitive participants, the effect of Familiarity was again significant, *F* (1, 16) = 9.44, *p* = .007, with a large effect size, 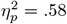, 95% CI [0.28, 1.00]. Estimated marginal means indicated a more negative N400 response to familiar sequences (*M* = *−*4.30, *SE* = 1.01, 95% CI [*−*6.44, *−*2.17]) than to unfamiliar sequences (*M* = *−*1.13, *SE* = 0.53, 95% CI [*−*2.26, 0.00]), *t*(16) = *−*3.07, *p* = .007, difference = *−*3.17, *SE* = 1.03. For insensitive participants, the effect of Familiarity was also robust, *F* (1, 49) = 20.11, *p <* .001, with a large effect size, 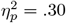, 95% CI [0.14, 1.00]. The N400 was more negative for familiar items (*M* = *−*2.80, *SE* = 0.48, 95% CI [*−*3.77, *−*1.84]) than for unfamiliar items (*M* = *−*0.66, *SE* = 0.25, 95% CI [*−*1.15, *−*0.17]), *t*(49) = *−*4.48, *p <* .001, difference = *−*2.14, *SE* = 0.48.

Together, these analyses demonstrate robust neural sensitivity to familiar shape sequences in both learner groups. Despite the absence of behavioral evidence of learning in the Insensitive group, their ERP responses showed significant familiarity effects, suggesting that both N100 and N400 components are capable of indexing learning that may not be expressed behaviorally.

### Model with Sensitiviy as a continous variable

To verify that the null Sensitivity effect was not an artifact of dichotomizing the sample, we re-estimated the N100 model using each participant’s z-scored *d*′ as a continuous predictor. The pattern of results remained unchanged. Familiarity showed a robust main effect, *F* (1, 65) = 29.49, *p <* .001, partial 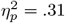, with more negative amplitudes for familiar than unfamiliar sequences (*M* = *−*3.18 *µ*V, *SE* = 0.44, 95% CI [–4.06, –2.30] vs. *M* = *−*0.78 *µ*V, *SE* = 0.23, 95% CI [–1.23, –0.32]). Neither the main effect of *d*′ (*F* (1, 65) = 2.04, *p* = .158, partial 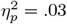) nor the *d*′*×* Familiarity interaction (*F* (1, 65) = 1.36, *p* = .249, partial 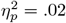) reached significance. The fixed-effect variance explained by the model remained modest 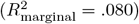, with *d*′ accounting for only about 3% of the explainable variance. These findings reinforce the conclusion that N100 amplitude tracks structural familiarity across participants but does not scale with behavioral sensitivity. These results are consistent with the findings from the categorical Sensitivity analysis.

We conducted the same analysis for the N400 time window, again using participants’ z-scored *d*′ scores as continuous predictors. As in the N100, Familiarity showed a strong main effect, *F* (1, 65) = 36.37, *p <* .001, partial 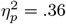, with more negative amplitudes for familiar than unfamiliar sequences (*M* = *−*2.00 *µ*V, *SE* = 0.55, 95% CI [–3.10 –0.90] vs. *M* = 1.77 *µ*V, *SE* = 0.50, 95% CI [0.76, 2.77]). Neither the main effect of *d*′ (*F* (1, 65) = 0.26, *p* = .614, partial 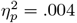) nor the interaction with Familiarity (*F* (1, 65) = 0.22, *p* = .640, partial 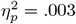) was significant. The marginal *R*^2^ was again low 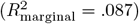, indicating that individual differences in behavioral sensitivity explained little additional variance in N400 amplitude.

Together, these results confirm that both the N100 and N400 reliably index structural familiarity in the visual stream. However, neither component shows graded sensitivity to behavioral performance as measured by *d*′, suggesting that the underlying learning mechanisms reflected in these ERP components operate largely independently of explicit behavioral discrimination.

### Model including Accuracy

While our primary analyses omitted Accuracy due to unbalanced trial counts, accuracy data can nevertheless clarify when and how strongly neural responses align with behavioral performance. Specifically, including Accuracy allows us to assess whether neural sensitivity to statistical structure varies systematically between correct and incorrect trials. To test this, we extended our models to include Accuracy (correct vs. incorrect) as an additional fixed effect, along with Familiarity and Sensitivity. The model also included by-subject random slopes for both Familiarity and Accuracy, and a random intercept for each electrode nested within subject.

In the N100 time window, we again observed a significant main effect of Familiarity, *F* (1, 65) = 27.44, *p <* .001, partial 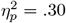, with more negative amplitudes for familiar than unfamiliar sequences (*M* = *−*3.55 *µ*V, *SE* = 0.51, 95% CI [–4.56, –2.54] vs. *M* = *−*0.90 *µ*V, *SE* = 0.26, 95% CI [–1.42, –0.37]). The main effect of Sensitivity was not significant, *F* (1, 65) = 2.50, *p* = .119, and no main effect of Accuracy was observed, *F* (1, 65) = 0.22, *p* = .644. No two-way interactions reached significance, but a significant three-way interaction emerged among Familiarity, Sensitivity, and Accuracy, *F* (1, 1673) = 56.01, *p <* .001, partial 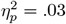. The model explained 9.0% of variance via fixed effects 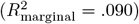 and 71.6% overall 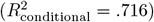.

Post hoc comparisons revealed a crossover pattern. For sensitive participants, the difference in amplitude between familiar and unfamiliar sequences was larger on correct trials (*M*_diff_ = *−*4.36, *SE* = 0.90; *d* = *−*0.82, *p <* .0001) than on incorrect trials (*M*_diff_ = *−*2.66, *SE* = 0.52; *d* = *−*0.43, *p* = .12). In contrast, for insensitive participants, the difference in amplitude between familiar and unfamiliar sequences was larger on incorrect trials (*M*_diff_ = *−*2.66, *SE* = 0.52; *d* = *−*0.65, *p <* .0001) than on correct trials (*M*_diff_ = *−*1.62, *SE* = 0.52; *d* = *−*0.40, *p* = .011). Thus, N100 amplitude differences tracked behavioral performance for sensitive participants but diverged from accuracy in the insensitive group.

In the N400 time window, a similar pattern emerged. Familiarity again showed a robust main effect, *F* (1, 65) = 30.88, *p <* .001, partial 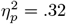, with more negative amplitudes for familiar than unfamiliar sequences (*M* = *−*1.92 *µ*V, *SE* = 0.63, 95% CI [–3.19, –0.66] vs. *M* = 2.06 *µ*V, *SE* = 0.58, 95% CI [0.90, 3.21]). The main effects of Accuracy, *F* (1, 65) = 1.13, *p* = .291, and Sensitivity, *F* (1, 65) = 0.57, *p* = .453, were not significant. However, as in the N100 analysis, a significant three-way interaction again emerged among Familiarity, Sensitivity, and Accuracy, *F* (1, 1673) = 60.36, *p <* .001, partial 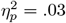. The model explained 9.7% of variance via fixed effects 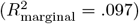 and 72.8% overall 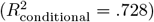.

Follow-up comparisons showed that sensitive participants exhibited a strong familiarity effect on correct trials (*M*_diff_ = *−*5.99, *SE* = 1.27; *d* = *−*0.91, *p <* .0001), but not on incorrect trials (*M*_diff_ = *−*2.82, *SE* = 0.27; *d* = *−*0.36, *p* = .12). Insensitive participants, by contrast, showed stronger effects on incorrect trials (*M*_diff_ = *−*4.41, *SE* = 0.74; *d* = *−*0.79, *p <* .0001) than on correct trials (*M*_diff_ = *−*2.69, *SE* = 0.74; *d* = *−*0.47, *p* = .002). These results indicate that ERP sensitivity to statistical structure in the N400 window is behaviorally aligned for sensitive participants, but dissociated from accuracy for the insensitive group. (See Fig 2)

**Fig 2.**
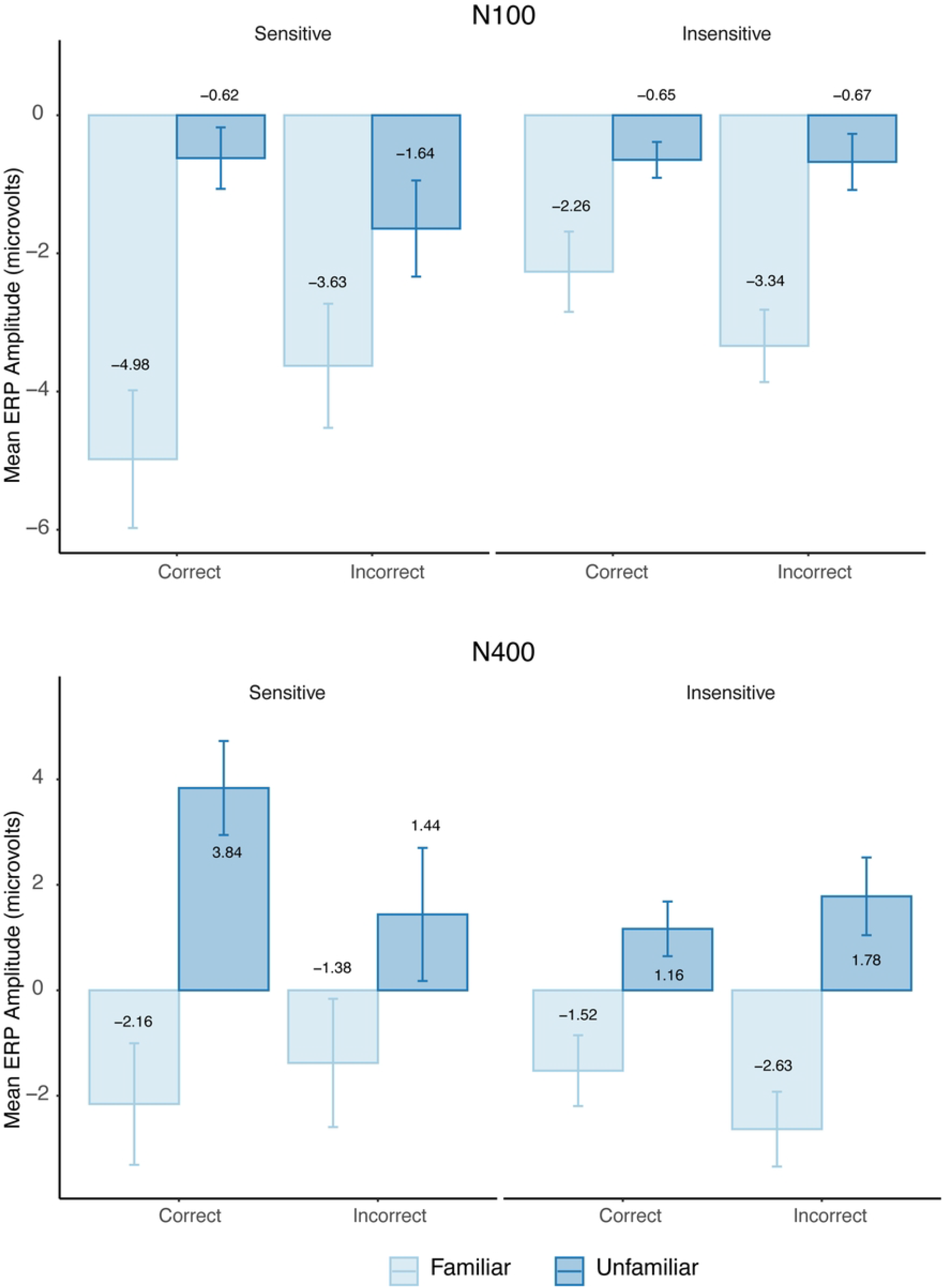
Bar plots showing the three-way interaction between Sensitivity, Familiarity, and Accuracy. Mean ERP amplitudes are plotted for the N100 (top panel) and N400 (bottom panel) time windows. Each panel displays bar plots grouped by Sensitivity (Sensitive vs. Insensitive), with bars representing familiar and unfamiliar sequences, separated by trial Accuracy (Correct vs. Incorrect). In the N100 window, sensitive participants show a larger familiarity effect (difference between familiar and unfamiliar amplitudes) on correct trials, whereas insensitive participants show a larger effect on incorrect trials. A similar pattern is observed in the N400 window, where the familiarity effect is stronger on correct trials for sensitive participants but stronger on incorrect trials for insensitive participants. Error bars reflect standard error of the mean.

Together, these findings suggest that while N100 and N400 amplitudes reliably differentiate previously encountered (familiar) shape sequences from novel ones, the relationship between neural and behavioral measures of learning differs across individuals. In sensitive participants, neural responses tracked overt task performance, consistent with explicit awareness of the learned regularities or stronger task engagement. In contrast, insensitive participants showed neural differentiation between familiar and unfamiliar sequences despite performing at or below chance behaviorally—suggesting the presence of implicit learning not accessible to conscious report, a finding that underscores the utility of ERPs for detecting learning-related brain activity that may not be reflected in overt behavior.

For full transparency, complete model outputs—including fixed effects, estimated marginal means (EMMs), and effect size estimates—are provided in the Supporting Information (see Summary of linear mixed-effects model results).

## Discussion

This study investigated how neural responses to statistical structure in visual input vary as a function of prior exposure to shape sequences and individual differences in behavioral performance. Using event-related potentials (ERPs), we focused on two components—the N100 and N400—that have previously been associated with early attentional orienting and later contextual integration, respectively. Our investigation was guided by two central questions: First, do the N100 and N400 components reliably distinguish between shape sequences that were previously encountered during familiarization and those that were novel, indicating neural sensitivity to learned structure? Second, do these neural responses vary systematically with individual differences in behavioral performance on a post-exposure familiarity judgment task? Our findings provide clear answers to both questions and offer new insights into the neural dynamics of statistical learning across individuals.

Both the N100 and N400 components showed robust modulation by Familiarity, with significantly more negative amplitudes for unfamiliar compared to familiar sequences. These results confirm that SL influences neural responses at both early perceptual (N100) and later integrative (N400) stages of processing. They align with prior work in auditory SL and extend it to the visual modality using a well-powered sample, with substantial effect sizes for the main effect of Familiarity in both components (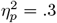 for N100; 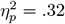 for N400).

However, one of the most striking findings of this study was that statistically familiar sequences elicited significant ERP effects—not only in sensitive participants who demonstrated above-chance behavioral performance, but also in insensitive participants whose responses failed to reliably distinguish familiar from unfamiliar items. These results contrast with findings from several auditory studies in which high-performing learners often exhibit stronger ERP responses to statistically structured sequences [11, 12].

Moreover, the relationship between these ERP effects and the overt responses differed by group: for sensitive participants, familiarity effects were largest on correct trials—when they accepted familiar triplets or rejected unfamiliar ones—indicating that ERP responses aligned with successful behavioral discrimination. In contrast, for insensitive participants, the largest familiarity effects emerged on incorrect trials—when they rejected familiar sequences or accepted unfamiliar ones—suggesting a disconnect between neural sensitivity and overt task performance. This crossover interaction points to a critical distinction in how learned structure is represented and accessed across individuals and raises broader questions about the relationship between implicit learning, conscious awareness, and behavioral expression.

The present discussion is organized around three central questions that emerged from our findings. First, we examine the surprising observation that participants classified as behaviorally insensitive nevertheless exhibited clear neural responses to statistical regularities, exploring possible explanations for this neural–behavioral dissociation. Second, we consider the conditions under which N100 effects arise in statistical learning tasks, focusing on why the current study revealed robust N100 modulation where previous visual SL studies did not. Finally, we interpret the three-way interaction among familiarity, correctness, and sensitivity, weighing alternative accounts of this finding and its implications for understanding how implicit learning signals are accessed and used during decision-making.

### Neural Evidence of Learning in the Absence of Behavioral Sensitivity

The robust N400 familiarity effects that we observed in both sensitive and insensitive participants challenge the common assumption that neural markers of learning directly mirror behavioral performance. These findings are particularly striking given that insensitive participants, by definition, showed little or no behavioral evidence of having acquired the statistical structure embedded in the sequences. How, then, can we account for the presence of reliable N400 modulation in these individuals?

We propose that this apparent discrepancy reflects differences in the time course of learning, rather than differences in whether learning occurred. Specifically, neural sensitivity to statistical structure—as indexed by the N400—may emerge earlier, and more gradually, than overt behavioral sensitivity. This proposal is supported by longitudinal evidence from prior studies. In a foundational auditory SL study, Abla et al. [12] recorded ERPs during the exposure phase and classified participants into high, middle, and low behavioral performers. They observed that high-performing learners exhibited strong N400 responses early in exposure, which gradually declined over time—presumably reflecting consolidation and reduced processing demands for familiar structures. In contrast, middle-performing learners showed a delayed increase in N400 amplitude across exposure blocks, while low performers showed little or no N400 modulation.

This trajectory is consistent with an inverted-U pattern in the N400 component: N400 amplitude increases as structure is initially detected and integrated into a developing representational system, then declines as the structure becomes familiar, stable, and more easily processed. A similar pattern was reported by Cunillera et al. [17], who observed that although N400 amplitude decreased in later blocks of a speech segmentation task, it remained robustly above baseline—indicating sustained sensitivity to the newly acquired structure even after repeated exposure.

In this framework, the N400 reflects the brain’s effort to integrate the current input into a broader internal model of learned structure. When structure is entirely absent, as in random or unsegmented input, there is no basis for prediction or integration, and thus the N400 is minimal. As structure emerges, the developing representations place increasing demands on integration processes, and the effort required to incorporate less stable or less familiar sequences into an internal model results in large N400 responses. With continued exposure and consolidation, familiar structures become easier to process, requiring less neural effort, and hence resulting in a smaller N400.

This view aligns well with classical findings from lexical-semantic ERP research. In traditional lexical decision tasks, pronounceable nonwords typically elicit the largest N400s, followed by real words, and then consonant strings, which elicit minimal or no N400 at all [10]. This hierarchy reflects a gradient of processing difficulty and integration effort. Nonwords are word-like enough to activate the lexical system but lack stable representations, leading to heightened neural processing. Real words, by contrast, are familiar and richly connected in semantic memory, requiring less effort to integrate. Consonant strings are not viable lexical candidates and thus evoke little engagement.

A similar gradient appears to operate in SL paradigms. Random sequences are like consonant strings: unstructured and unprocessable, they evoke minimal N400 responses. Newly learned but fragile structural units are like pronounceable nonwords: they are processed as candidates for integration but lack firm anchoring in the learner’s representational system, generating large N400 responses. Fully familiarized structures resemble real words: they are easily integrated and elicit smaller N400s. Thus, while SL paradigms lack explicit semantic content, the functional demands placed on the learner’s system—namely, the need to integrate current input into a probabilistic model of structure—are conceptually analogous to lexical-semantic integration.

In the present study, we recorded ERPs not during the exposure phase, but during a post-familiarization test phase. By this point, sensitive learners may have consolidated the structure to the point where it is readily integrated, leading to moderate N400 amplitudes. In contrast, insensitive learners—whose neural trajectories may lag behind—may be in the early-to-mid phase of the inverted-U pattern, where integration is still effortful but structure has become internally coherent enough to support prediction. Thus, we observe robust N400 effects in these individuals, even though their overt behavior has not yet caught up.

This account is consistent with findings from Sanders and Neville [15], who reported larger N400 amplitudes for structured words than for unstructured nonwords in an auditory SL paradigm—a reversal of classical N400 effects that they described as “unexpected.” While atypical in the context of semantic processing, such findings make sense in light of the inverted-U trajectory of structure learning: novel but coherent sequences are harder to integrate than familiar or meaningless ones. More generally, these results highlight the N400’s sensitivity not just to semantic incongruity but to representational instability—the difficulty of integrating a unit that is being learned, rather than one that is already known.

Finally, our use of linear mixed-effects models likely increased our sensitivity to these subtle neural signals. Unlike traditional averaging methods, mixed models can account for both subject- and item-level variability, making them better suited to detect effects that are weak or inconsistent across trials. This may explain why prior studies failed to observe N400 effects in low-performing learners, even if those effects were present at a neural level.

In sum, the N400 familiarity effects observed in insensitive participants should not be taken as anomalous or contradictory. Rather, they reflect a coherent and developmentally plausible pattern: neural learning precedes behavioral learning. The N400, in this context, marks the brain’s sensitivity to regularities that have been implicitly acquired but not yet behaviorally expressed. This interpretation helps to reconcile diverse findings in the SL literature and underscores the value of ERPs as a window into latent learning processes that might otherwise go undetected.

### When Do N100 Effects Arise in Statistical Learning?

Prior ERP studies of statistical learning (SL) have produced mixed results regarding N100 modulation. In some auditory paradigms, N100 enhancement has been reported at word or triplet onsets, particularly when external segmentation cues such as prosodic stress [15, 16] or very strong transitional probability (TP) manipulations [13, 14] are present. These cues are believed to direct attention to likely boundaries, enhancing early perceptual processing and increasing N100 amplitude. For example, Cunillera et al. showed that stressed syllables elicited larger N1 and N400 responses, while Soares and colleagues observed stronger N100 and N400 responses to high-TP items. However, in the absence of such cues—such as when only moderately strong TPs signal structure—N100 effects are often absent. For instance, Abla and Okanoya [18] reported clear N400 effects but no N100 modulation in a visual SL task with no explicit boundary markers.

We see two possible explanations. The first concerns the timing of ERP measurement. In our study, ERPs were recorded during a post-exposure test phase, after participants had the opportunity to internalize the statistical structure of the shape sequences. In contrast, prior studies such as that of Abla and Okanoya [18] recorded ERPs during the exposure phase, when structure-driven processing may not yet have emerged. If the N100 reflects attention guided by learned structure—as we propose—then such effects may only become detectable after learning has occurred and when that knowledge can be prospectively deployed. This distinction between online learning and offline recognition may account for the divergent N100 findings across studies.

The second possibility relates to how the N100 was measured. As noted above, in our study, we measured ERPs during the test phase where sequences were presented in isolation, with clearly marked boundaries around each triplet. In contrast, Abla and colleagues recorded ERPs during the familiarization phrase in which triplets were presented as a continuous stream of unsegmented input. If the N100 reflects a perceptual response to the onset of a newly learned unit, it may be more easily evoked when that onset is unambiguous—as in isolated trials—than in continuous streams, where the beginning of each unit must be inferred. On this view, the presence of N100 modulation in our study may reflect not only learned structure but also the clarity of item segmentation at test.

Supporting this view, we observed reliable N100 modulation even in the absence of overt segmentation cues, and across both sensitive and insensitive participants. This suggests that the N100 in our task does not reflect purely exogenous cue-based processing, but rather endogenously guided attention shaped by the statistical structure of the input. Once statistical structure has been learned, it may act as a cue in its own right—driving early perceptual grouping and enhancing N100 amplitude. Prior studies that failed to observe N100 effects may not have provided sufficient learning opportunities, or may have captured neural responses too early in the learning trajectory.

### Interpreting the Three-Way Interaction of Familiarity, Sensitivity, and Accuracy

Our analyses revealed a significant three-way interaction among Familiarity, Accuracy, and Sensitivity in both the N100 and N400 components. For sensitive participants, the ERP familiarity effect—defined as the difference between responses to familiar and unfamiliar triplets—was larger on correct trials than on incorrect ones. For insensitive participants, this pattern was reversed: familiarity effects were larger on incorrect trials. As this analysis was exploratory and not guided by a priori hypotheses, the results should be interpreted with caution. Nevertheless, following standard logic for interpreting correlational findings, we consider three broad classes of explanation:

#### Accuracy influences ERP responses

In this account, the N100 and N400 reflect neural responses that may be amplified when there is dissonance between the brain’s implicit familiarity signals and the participant’s subsequent behavioral decision—a form of prediction–response conflict. That is, when the brain registers a stimulus as familiar but the participant responds otherwise, the resulting mismatch may enhance ERP amplitudes, potentially reflecting internal error monitoring or conflict detection. However, this explanation appears implausible given the temporal structure of the task. ERPs were measured within the first 400 ms following the onset of the initial shape, while the behavioral decision occurred approximately 2000 ms later. Thus, this account stretches temporal logic and assumes that neural responses can be shaped by events that have not yet occurred.

#### ERP responses influence behavioral accuracy

On this view, for both sensitive and insensitive participants, the brain showed clear evidence of detecting statistical familiarity, as reflected in larger N400 responses to familiar sequences than to unfamiliar ones. However, only sensitive participants appeared to use this neural familiarity signal to guide their decisions. When the N400 signal was strong—indicating that the brain recognized a familiar pattern—these participants were more likely to respond “yes,” resulting in high accuracy. Thus, the N400 familiarity effect was larger on correct trials than incorrect ones.

In contrast, insensitive participants often failed to align their behavioral responses with the neural familiarity signal. On trials where the N400 was large—indicating that the brain registered the sequence as familiar—they frequently responded “no,” producing incorrect judgments (misses). This pattern suggests that they either misinterpreted the signal (e.g., treating heightened neural activity as a marker of novelty or uncertainty) or disregarded it altogether. On correct trials, by contrast, the N400 signal tended to be weaker, and responses may have been more variable or based on other factors such as guessing or low-level cues. As a result, the familiarity effect in this group was larger on incorrect trials than on correct ones.

A similar pattern was observed for the N100, suggesting that early perceptual responses were also modulated by familiarity and linked to decision outcomes. On this hypothesis, the correlation between the N100 and behavior may reflect participants’ use of early perceptual cues to guide their decisions—or it may indicate that the N100 influenced later processing stages, including the N400.

#### A third variable modulates both ERP and behavioral responses

While the possibility of a third variable cannot be ruled out, it is difficult to identify a plausible candidate variable that would modulate both ERP amplitudes and response accuracy in opposite directions for the two groups. Factors such as attention, motivation, or fatigue might influence overall performance, but are unlikely to produce the specific interaction pattern observed here unless they systematically affect signal interpretation differently in each group.

In sum, the observed three-way interaction underscores the importance of distinguishing between neural sensitivity to statistical regularities and the effective use of that information in behavior. While both groups demonstrated robust neural differentiation between familiar and unfamiliar sequences, only sensitive participants consistently aligned their behavioral responses with this signal. These findings highlight the value of ERP measures in uncovering latent learning processes that may not be evident in overt behavior, and they point to meaningful individual differences in how learned structure is accessed and applied during decision-making.

## Conclusion

This study provides clear evidence that visual statistical learning modulates neural responses at both early (N100) and late (N400) stages of processing. We can state with confidence that both sensitive and insensitive participants exhibited robust ERP differentiation between familiar and unfamiliar shape sequences, demonstrating that the brain is capable of tracking statistical structure even in the absence of overt behavioral learning. These effects were observed using a well-powered sample, consistent analytical procedures, and convergent results across components, strengthening the reliability of this finding.

More tentatively, our exploratory analyses suggest that the relationship between neural sensitivity and behavior may differ across individuals. Sensitive participants appeared to align their decisions with the strength of the neural familiarity signal, while insensitive participants often failed to do so—possibly due to differences in signal interpretation, access, or response strategy. Although the pattern is compelling, the mechanisms underlying this dissociation remain speculative and warrant further investigation. Future studies with longitudinal designs or real-time neural decoding could clarify how implicit learning signals are accessed and used—or misused—during decision-making.

Together, these findings support the broader conclusion that ERPs provide a valuable window into latent learning processes and individual differences in their behavioral expression. They underscore the importance of considering both neural and behavioral measures when characterizing learning, particularly in domains where implicit knowledge may not be readily accessible through overt performance.

## Supporting information

**Table S1. Full model outputs for N100 and N400 ERP analyses.** This table presents summary statistics for four linear mixed-effects models for each component (N100 and N400). It includes fixed effects, F-statistics, p-values, partial eta-squared effect sizes, marginal R^2^, estimated marginal means (EMMs), standard errors, and Cohen’s d with 95% confidence intervals for the familiarity effect across sensitivity conditions.

**Table 1.**
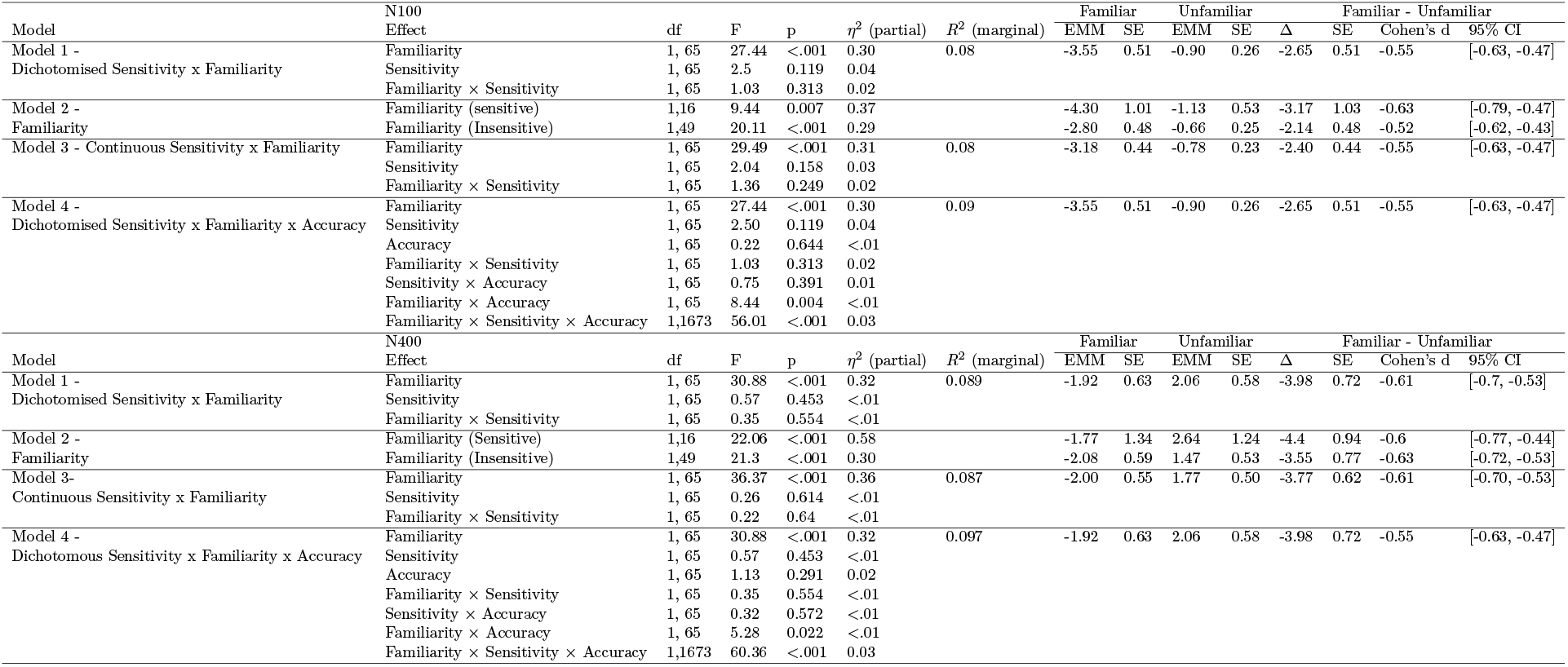
Summary of linear mixed-effects model results.

## Acknowledgments

This research was initiated at Hampshire College during Joanna Morris’s appointment there and completed at Providence College, where data analysis and manuscript preparation were conducted.

Research reported in this manuscript was supported by the Rhode Island IDeA Network of Biomedical Research Excellence from the National Institute of General Medical Sciences of the National Institutes of Health under grant number P20GM103430. Additional support was provided by the Center for Engaged Learning at Providence College, which funded Joemari Pulido, Tess Rooney, Natalia Alzate, Ethan Moore, and Crismar Ramos-Marte to work on the project over several summers. The research also benefited from multiple Faculty Development Grants awarded to Joanna Morris during her tenure at Hampshire College.

We are especially grateful to Ani Alaberkyan, who served as a full-time research assistant and played a central role in running the lab for a year. We also thank the following individuals affiliated with the Department of Psychology at Providence College and the School of Cognitive Science at Hampshire College for their valuable contributions to experiment setup and data collection: Kirsten Lydic, Sophie Fulghum, Dylan Palumbo, Maeve Garrity, Aleksandra Kurylowicz, Alexandra Saneff, Sarah Debian, Kenny Diaz, Amelia Bycoff, Abbie Okon and Matthew Dunn

